# The virome of *Drosophila suzukii*, an invasive pest of soft fruit

**DOI:** 10.1101/190322

**Authors:** Nathan C. Medd, Simon Fellous, Fergal M. Waldron, Anne Xuéreb, Madoka Nakai, Jerry V. Cross, Darren J. Obbard

## Abstract

*Drosophila suzukii* (Matsumura) is one of the most damaging and costly pests to invade temperate horticultural regions in recent history. Conventional control of this pest is challenging, and an environmentally benign microbial biopesticide is highly desirable. A thorough exploration of the pathogens infecting this pest is not only the first step on the road to the development of an effective biopesticide, but also provides a valuable comparative dataset for the study of viruses in the model family *Drosophilidae.* Here we use a metatransciptomic approach to identify viruses infecting this fly in both its native (Japanese) and invasive (British and French) ranges. We describe 18 new RNA viruses, including members of the Picornavirales, Mononegavirales, Bunyavirales, Chuviruses, *Nodaviridae, Tombusviridae, Reoviridae,* and Nidovirales, and discuss their phylogenetic relationships with previously known viruses. We also detect 18 previously described viruses of other *Drosophila* species that appear to be associated with *D. suzukii* in the wild.

## Introduction

*Drosophila suzukii* (Matsamura) is an invasive dipteran pest of soft fruit belonging to the subgenus Sophophora. Unusual within the genus, the larvae are well adapted to feeding on ripe fruit still on the plant, adult females possess a heavily sclerotized saw-like ovipositor that allows oviposition under the skin of ripening fruit, and their olfactory system is adapted to respond to fruit rather than microbe volatiles (Karageorgi et al., 2017). These evolutionary innovations may aid the establishment of this species in novel habitats across the globe (Atallah et al., 2014, Poyet et al., 2015).

First described in Japan in 1916 (Kanzawa, 1935, Matsumura, 1931), *D. suzukii* was reported to be widely distributed in Japan shortly after (Kanzawa, 1939). It was recorded across Asia during the last century (Peng, 1937, Kang and Moon, 1968, Parshad and Duggal, 1965, Okada, 1976, Toda, 1991, Sidorenko, 1992, Amin ud Din et al., 2005), with the first records outside of Asia coming from Hawaii in the 1980′s (Kaneshiro, 1983). Since its detection in 2008 in the southern states of the USA (Bolda, 2008) and Spain (Calabria et al., 2012), *D. suzukii* has spread northwards, and was recorded for the first time in the UK in 2012 (Harris and Shaw, 2014). Records now stretch from Sweden (Manduric, 2017) to Argentina (Lue et al., 2017), with secondary invasions thought to be responsible for populations detected in South America and the Indian Ocean Islands (Fraimout et al., 2017).

The damage *D. suzukii* has caused in the fruit growing regions of these countries has driven interest in many aspects of the pest′s biology, primarily to improve control methods (Asplen et al., 2015). Conventional chemical control of *D. suzukii* is challenging because oviposition occurs so close to harvest that management of pesticide residues during crop treatment is of concern (Swoboda-Bhattarai and Burrack, 2015). *D. suzukii* also has a broad host range allowing it to exploit natural refugia, including many wild *Prunus* and *Rubus* spp. (Walsh et al., 2011, Cini et al., 2012, Mitsui et al., 2010, Poyet et al., 2014). An effective biological control agent of *D. suzukii*, compatible with integrated management techniques (Stern et al., 1959), would therefore be highly desirable to horticulturalists worldwide.

Entomopathogenic viruses have the potential for use as environmentally benign, species-specific biological control agents, with certain groups of viruses being used to effectively control insect pests in a range of settings (Hunter-Fujita et al., 1998). The most successful viral control agents to-date are members of the *Baculoviridae,* with the nuclear polyhedrosis viruses (NPVs) and granulosis viruses (GVs) finding commercial success against lepidopteran pests in forestry and orchard crops, respectively. These viruses produce polyhedrin occlusion bodies that encase infectious virions during the dispersal stage of the viruses′ lifecycle. These protein occlusions protect the virus from environmental degradation, and prolong infectivity in the environment (Bishop et al., 1988, Elgee, 1975). For this reason these viruses have been the focus of viral biopesticide development since their first commercial use in 1975 (Shieh and Bohmfalk, 1980). However, despite the relative success of the *Baculoviridae,* other viral taxa have also been advocated for control purposes. For example: members of the *Nudiviridae,* for use against Rhinoceros beetle (Huger, 2005); a member of the *Reoviridae* for use against Masson pine moth (Peng et al., 2000), and certain viruses of the *Parvoviridae* for use against a range of pests (Bergoin and Tijssen, 1998). All have shown promise as control agents, despite not achieving commercial success.

As well as identifying some of the natural enemies that could be harnessed to control *D. suzukii* populations, understanding the nature of viral infections in this species may help us better understand the reasons for its geographical spread and establishment (Mitchell and Power, 2003, Torchin et al., 2003, Colautti et al., 2004). In particular, the changing pathogen environment that invasive species encounter in the process of invasion is of interest to the field of ecological immunology, in that relative immune investment is predicted to depend upon the diversity of pathogens experienced its new range (Joshi and Vrieling, 2005, Colautti et al., 2004, Blossey and Notzold, 1995). However, thorough surveys of pathogen diversity in wild invaders remain relatively rare (but see Liu and Stirling, 2006). The genus *Drosophila* is one of the few invertebrate genera in which wild viral pathogen diversity has been explored, with recent virus discovery studies in the genus describing over 50 new viruses (Webster et al., 2016, Webster et al., 2015). Furthermore, a history of intensive investigation of the antiviral immunity of *D. melanogaster* (Xu and Cherry, 2014, Sabin et al., 2010, Huszar and Imler, 2008, Kemp and Imler, 2009, Mussabekova et al., 2017, Bronkhorst and van Rij, 2014, Zambon et al., 2006), means that the viruses of *D. suzukii* may provide a valuable comparative system for the study of immune system evolution.

Here we report the results of a metatransciptomic survey of virus-like sequences associated with *D. suzukii* in both its native (Japanese) and invasive (British and French) ranges. We describe 18 new RNA viruses, representing 10 different virus families, and confirm their presence in RNA pools using RT-PCR. We place these viruses in the phylogenetic context of recent metatransciptomic studies in the host genus (Webster et al., 2015, Webster et al., 2016) and in invertebrates as a whole (Shi et al., 2016).

## Methods

### Sample collection

We collected 4450 individual *D. suzukii* across a three-year period between September 2013 and September 2016, including 230 larvae in 2016. We initially focussed on flies in their European invasive range, with sampling subsequently extended to include surveys of flies from native SE Asian range. Flies were collected near Montpellier, France (43.59 N, 3.78 E) in 2013 (collection by AX and SF); in Kent, UK (51.284 N, 0.465 E) during the late summer of 2014, 2015 and 2016 (NCM); and in three locations across Honshu, Japan, during May 2016 (NCM and MN): Tokyo University of Agriculture and Technology, Fuchu (35.683 N, 139.481 E); Naganuma Park, Tokyo (35.637 N, 139.375 E); Shimaminami Shima, Yamagata Prefecture (38.351 N, 140.276 E); Agriculture Total Centre Kaju Research Institute, Fukushima (37.813 N, 140.443 E); and Fuefukigawa Fruit Park, Yamanashi (35.700 N, 138.666 E). We used a combination of commercial bait traps with cotton soaked in a proprietary liquid attractant (DROSO TRAP^®^ and DROS′ATTRACT^®^, Biobest, Belgium, NV), and a standard sweep net to catch adult flies. Traps, hung at field margin and woodland sites, were collected at intervals of two to three days. All individuals were sorted into vials by trap and species within three hours of collection. We aimed to morphologically identify all species of *Drosophila* caught (Bächli et al., 2004), however, we also subsequently examined RNA pools for potential contamination due to misidentification. Other species of *Drosophila* were caught in these traps and we collected them together with *D. suzukii,* but they were not analysed further. Wild-collected flies were maintained on solid agar/sugar medium, before being macerated in sterile Ringer′s solution (to allow for future experimental virus culture and isolation). In addition to adult fly samples larvae were extracted from infested fruit collected in 2016 from UK and Japan with sterile forceps. Although no *Drosophila* pathogens have previously been reported from the larval stage alone, through their collection we aimed to address the possibility that our sampling method was biased towards mobile adult flies able to respond to attraction based traps.

We pooled trap catches from within a sampling location and immediately extracted RNA from a subsample of the fly (or larva) homogenate using TRIzol® (Invitrogen), before storage at −80°C. We treated pooled RNA samples for possible DNA contamination using DNase (Turbo DNA-free, Ambion) prior to library preparation. To verify RNA quality we tested for contamination using Qubit® and Nanodrop® spectrophotometers. For flies collected in the UK and Japan, library preparation and strand specific sequencing was performed by Edinburgh Genomics (Edinburgh, UK) using Illumina NGS library preparation kits and the Illumina Hi-Seq platform with 120 or 150nt paired end reads. To increase representation of viral and host protein coding RNAs, all libraries underwent depletion of rRNA using Ribo-Zero rRNA Removal Kit (Illumina). Flies collected in France during 2013 were sequenced separately at Beijing Genomics Institute (BGI tech solutions, Hong Kong) using paired-end 90nt reads using the HiSeq 2000 platform. These libraries underwent Duplex-Specific Thermostable Nuclease (DSN) normalisation and poly-A selection. This process, although enriching for viruses by rRNA depletion, biases virus discovery towards poly-adenylated genomic products only produced by certain viral taxa (e.g. Picornavirales). All raw reads have been submitted to the NCBI sequence read archive under project accession PRJNA402011 (Japan SRR6019484; France SRR6019487; Kent: SRR6019485, SRR6019486, and SRR6019488).

### Virus identification and Phylogenetic Analysis

To remove those reads derived from *Drosophila,* we mapped raw reads against the *D. suzukii* genome and transcriptome using Bowtie2 (Langmead and Salzberg, 2012) with the ′‐‐very-fast′ command-line option. We used Trimmomatic (Bolger et al., 2014) to quality trim and remove adapter sequences from the remaining unmapped raw reads (as pairs) using default parameters, before *de novo* assembly using Trinity version 2.2.0 (Grabherr et al., 2011), retaining a minimum contig length of 500nt. All raw unannotated contigs are provided in supporting file S1 (doi: 10.6084/m9.figshare.5649829). We concatenated all translations of all open reading frames (ORFs) in each resulting contig, and retained only those with an open reading frame of 150 codons or greater. These concatenated protein sequences were used to search against a custom database using Diamond (Buchfink et al., 2015) with a an e-value threshold of 0.01, retaining a single top hit. The target database comprised all of the viral proteins from the Genbank non-redundant protein database (‘nr’; Clark et al., 2016), and all of the prokaryote, protist, fungal, nematode, hymenopteran, and dipteran sequences from NCBI refseq protein database. Contigs for which the top hit was a virus were imported into Geneious®8.0.2 sequence analysis software (Kearse et al., 2012) for manual analysis. We grouped putative virus fragments taxonomically according to their initial best Diamond hit, assembled (Geneious) and manually curated them with reference to closest relatives in Genbank, to give the longest viral sequences consistent with the predicted protein content and structure of that virus taxon.

To infer phylogenetic relationships, we used RNA-dependent RNA polymerase (RdRp) coding sequences unless otherwise stated. The RdRp is generally the most conserved protein across RNA viruses, making it suitable for phylogenetic analysis of this diverse set of virus taxa (Koonin et al., 1993, Shi et al., 2016). RdRp gene sequences were translated and aligned with homologous sequences from their close relatives, as identified by BLAST (Altschul et al., 1990). To align multiple protein sequences we used ClustalW (Thompson et al., 2002) with BLOSSOM cost matrix (Henikoff and Henikoff, 1992). We manually identified regions of poor alignment at the 5′ and 3′ ends of the alignment and removed them before further analysis. All alignments are provided in supplementary material S2_Data (DOI: 10.6084/m9.figshare.5650117). We then inferred maximum-likelihood phylogenetic trees using PhyML 2.2.3 (Guindon and Gascuel, 2003) with the LG substitution model (Le and Gascuel, 2008). We calculated branch support using the Shimodaira-Hasegawa-like nonparametric version of an approximate likelihood ratio test implemented in PhyML (aLRT; Anisimova et al.,2011) (60). For clarity, the trees presented in figures 2, 4, and 6 are clades from within of larger trees (full trees provided in S3_Data, DOI: https://doi.org/10.6084/m9.figshare.5650132), realigned and reconstructed using the same methods.

### Detection by RT-PCR

To confirm the presence of the newly discovered viruses in original RNA pools we used Reverse Transcription PCR (RT-PCR) to screen for short amplicons of each virus′ longest ORF, where possible spanning part of the RdRp gene. We designed primers using the Primer3 (Rozen and Skaletsky, 1999) plugin for Geneious (Kearse et al., 2012), and where necessary manually adjusted oligoes to avoid polymorphic variants identified through pool sequencing, and to avoid synonymous positions at the 3’ end. RNA virus sequences identified by metagenomic methods may derive from viral elements endogenised into genomic DNA, if they are expressed (Katzourakis and Gifford, 2010). To test for endogenised viral elements (EVEs) we conducted PCRs (without a reverse transcription step) on nucleic acid samples that contained genomic DNA from the original phenol-chloroform extraction. As these RNA viruses do not produce a DNA intermediate, any viruses detectable by PCR in the DNA fraction are likely to be EVEs.

### Virus genome annotation

For viruses with complete, or near complete genomes, we were able to infer genome structure and identify protein functional domains by first identifying ORFs and then comparing these to the Conserved Domain Database with an expected value threshold of 5 × 10^−3^, and searching the NCBI ′nr′ protein database using BLASTp. Only ORFs of 100 amino acids or longer were annotated, unless notable similarity to closely related viruses was evident. ORFs of less than 200 amino acids that were nested completely with larger ORFs were disregarded, unless they displayed high similarity to known proteins.

### Distribution of RNA sequence reads across samples

To estimate the number of virus reads in each pooled sample, and to detect any cross-species contamination in fly collections, we mapped trimmed forward reads to all new and previously published *Drosophila* virus genomes (including multiple divergent isolates where they were available), a selection of *Drosophila* ribosomal sequences, and a short region of cytochrome oxidase 1 (COI) that has discriminatory power between *Drosophila* species. Sequences were mapped with Bowtie2 (Langmead and Salzberg, 2012) using the ′‐‐very-sensitive′ option. We report these after normalisation by the number of non-ribosomal reads and the length of each target sequence. We also apply an arbitrary lowest level detection threshold for each putative species of 0.5 total reads per Kb per million non-rRNA reads, to reduce spurious signals caused by low level species contamination, library barcode switching, and cross-mapping to close relatives.

## Results

In total, we generated approximately 280 million read pairs, ranging from 33 million pairs (UK – 2016) to 105 million pairs (France – 2013) per library. Our assemblies comprised between 18,431 (Japan – 2016) and 56,384 (UK – 2015) putative transcript contigs. Among these, we identified 18 new RNA viruses associated with *D. suzukii* (Table 1.). These viruses represent a variety of RNA virus taxa with positive sense single stranded (+ssRNA), negative sense single stranded (-ssRNA), and double stranded RNA (dsRNA) genomes, and include representatives of the Picornavirales, Mononegavirales, Bunyavirales, Chuviruses, *Nodaviridae, Tombusviridae, Reoviridae* and Nidovirales. We did not identify any DNA viruses despite active DNA virus infections being easily detected from RNA sequencing data. We do not report as new any viruses detected in *D. suzukii* that are identical, or near identical (>95% amino acid similarity in the polymerase), to previously published viruses. Those previously described viruses that were detected in *D. suzukii* are detailed in supplementary material (S1_table; DOI: 10.6084/m9.figshare.5650147) and relative read counts in each pool are shown in Figure 1.

**Table 1.** Novel viruses detected in *D. suzukii.* ^a^PCR reactions performed on cDNA. ^b^PCR reactions performed on extractions containing nuclear DNA.

| Provisional Name | Accession | Host | Taxon | Genome | Longest contig (kb) | Sample(s) | Detected by RT-PCR<sup>a</sup> | Detected by PCR<sup>b</sup> |
| --- | --- | --- | --- | --- | --- | --- | --- | --- |
| **Beult virus** | MF893261, MF893262 | Dsuz | Hepe-virga clade | +ssRNA | 12 | France2013, UK2014, UK2015, UK2016, Japan | + | - |
| **Saiwaicho virus** | MF893256 | Dsuz | Hepe-virga clade | +ssRNA | 10 | Japan2016 | + | - |
| **Luckshill virus** | MF893250 | Dsuz | Hepe-virga clade | +ssRNA | 3.5 | UK2016 | + | - |
| **Teise virus** | MF893259 | Dsuz | *Luteoviridae* | +ssRNA | 4.5 | France2013, UK2014, UK2015, UK2016, Japan2016 | + | - |
| **Tama virus** | MF893258 | Dsuz | *Sobemovirus* | +ssRNA | 3.5 | Japan2016 | + | - |
| **Medway virus** | MF893251 | Dsuz | *Sobemovirus* | +ssRNA | 2.7 | UK2014 | + | - |
| **Dsuz Nora virus** | MF893254 | Dsuz | *Picornaviridae* | +ssRNA | 12 | Japan2016 | + | - |
| **Naganuma virus** | MF893253 | Dsuz | *Nodaviridae* | +ssRNA | 1.6 | Japan2016 | + | - |
| **Fuefuki virus** | MF893247 | Dsuz | *Nidoviridae* | +ssRNA | 16 | Japan2016 | + | - |
| **Cyril virus** | MF893263 | Dsuz | Hepe-virga clade | +ssRNA | 1.5 | UK2016 | + | - |
| **Eccles Virus** | MF893265-MF893270 | Dsuz | *Reoviridae* | dsRNA | 4.2 | UK2014 | + | - |
| **Larkfield virus** | MF893249 | Dsuz | *Totiviridae* | dsRNA | 6 | UK2015 | + | - |
| **Snodland virus** | MF893257 | Dsuz | *Totiviridae* | dsRNA | 1.6 | UK2015 | + | - |
| **Mogami virus** | MF893252 | Dsuz | *Chuvirus* | -ssRNA | 10.5 | Japan2016 | + | - |
| **Ditton virus** | MF893264 | Dsuz | *Phasmaviridae* | -ssRNA | 7.3 | UK2015 | + | - |
| **Barming virus** | MF893260 | Dsuz | *Phleboviridae* | -ssRNA | 6.5 | UK2016 | + | - |
| **Notori virus** | MF893255 | Dsuz | *Phasmaviridae* | -ssRNA | 7 | Japan2016 | + | - |
| **Kiln Barn virus** | MF893248 | Dsuz | *Chuvirus* | -ssRNA | 3.7 | UK2016 | + | - |

**Figure 1.**
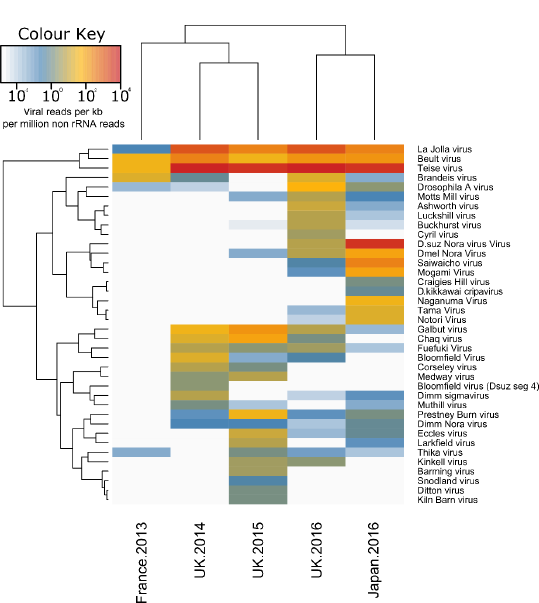
The heatmap shows the relative number of reads (log_10_ reads per kb per million non-ribosomal RNA reads) from each library mapping to each of the *Drosophila* viruses. Rows and columns are clustered by their similarity in read frequency on a log_10_ scale. A threshold for detection of 0.5 reads per kb per million non-rRNA reads was applied, however, a small amount of cross mapping is possible between closely related viruses and this may explain the detection of viruses with very low read counts. The low diversity of viruses in the France 2013 sample may be attributable to poly-A selection of RNA libraries. Created using the ′heatmap2’ function of the gplots package (Warnes et al., 2016) in R (R Core Team, 2017).

We have provisionally named these viruses according to the location from which the hosts were sampled. We have chosen not to include taxonomic or host information in the provisional name of the virus, as these are subject to change as phylogenetic relationships are revised and alternative or additional hosts discovered. The one exception to this rule is *D. suzukii* Nora Virus. This virus is sufficiently closely related to the *D. melanogaster* Nora virus and *D. immigrans* Nora virus that a name outside of this local scheme may cause confusion for future studies. During Phylogenetic analysis, a number of virus-like sequences were identified by BLAST in the public Transcriptome Shotgun Assembly Database (TSA). These have been included in analyses to help improve accuracy of phylogenetic inference, but are not further discussed.

### Viruses with single-stranded positive sense RNA genomes

Ten of the viruses described here are expected to encode their genomes in +ssRNA. Of these, Teise virus was found at the highest levels across samples. Teise virus is a sobemo-like virus closely related to Prestney Burn virus of *D. subobscura* (Webster et al., 2016) and Motts Mill virus of *D. melanogaster* (Webster et al., 2015), with 90.9% and 88.6% RdRp amino acid similarity respectively (Fig. 2, A). The single-stranded positive sense genome of these viruses comprises two unjoined fragments, which may represent subgenomic products (Webster et al., 2015, Shi et al., 2016, Tokarz et al., 2014) (Fig. 3), a structure consistent with its close relatives (Shi et al., 2016, Webster et al., 2015). Teise virus is the most geographically widespread virus of *D. suzukii,* with reads appearing in high numbers in both native and naturalised ranges (Fig. 1).

**Figure 2.**
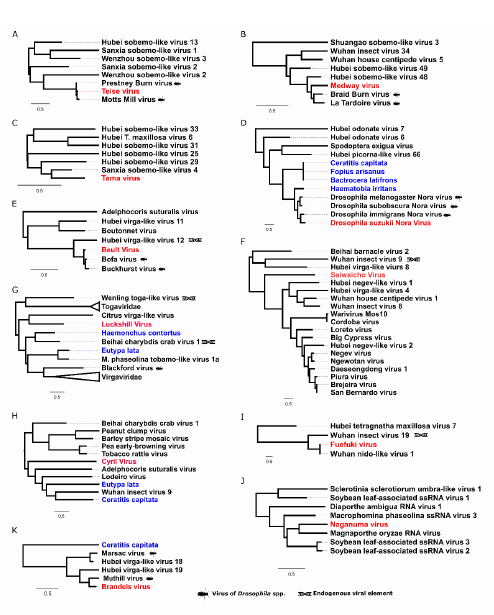
Positive sense single stranded RNA viruses. Midpoint-rooted, maximum-likelihood trees were inferred from viral polymerase or viral transferase (H only) sequences. Scale bar represents 0.5 substitutions per site. Putative viruses newly described in association with *D. suzukii* (red) are highlighted alongside virus-like sequences identified in public transcriptome datasets (blue). Viruses previously described as endogenous viral elements are also marked. Tree A,B,C: Sobemo-like viruses belonging to clusters within the Luteo-Sobemo clade; D: Noraviruses and related cluster of the Picora-Calici clade; E,F: Virga-like virus clusters nearby Cilevirus and Negeviruses in the Hepe-Virga clade; G: A small cluster of toga-like viruses neighbouring the Alphaviruses, *Togaviridae;* Hepe-virga clade; H: A cluster of Virga-like viruses constructed from transferase sequence; I: a cluster in the Nidoviruses close to the *Coronaviridae;.* J: Cluster of Nodaviruses within the Tombus-Noda clade; K: A cluster containing three *Drosophila* viruses within the Hepe-virga clade and distantly related to the *Virgaviridae* and *Bromoviridae.* Complete trees are provided in supporting file S3_data.

**Figure 3.**
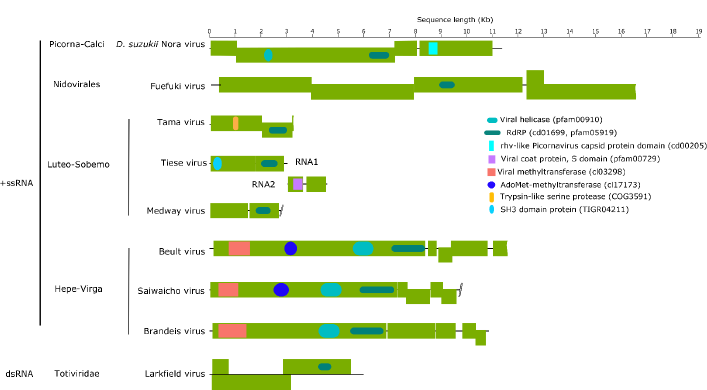
The structure of selected dsRNA and +ssRNA virus genomes for which we recover complete or near complete genome sequences. Outer (green) boxes represent boundaries of ORFs and inner boxes represent the relative position of conserved domains identified with reference to the NCBI Conserved Domain Database. Waved outer boxes represent incomplete ORFs and lines ending in slashes represent areas where genome is expected to contain further ORF not recovered from this analysis.

Medway virus (Fig. 2, B) shares close relationship to Braid Burn virus, previously described from *Drosophila subsilvestris* in the UK (Webster et al., 2016). These viruses belong to a clade of insect viruses distantly related to the Sobemo and Poleroviruses of plants (Shi et al., 2016). Medway virus appears at low copy-number in our samples with a small number of reads being detected in UK samples from 2014 and 2015. As for other viruses in this section of the Luteo-Sobemo group, the Medway virus genome probably consists of two genomic RNA segments. However, we were unable to detect the second RNA segment and we describe the virus only from an RNA fragment that contains two ORFs, including the RdRp (Fig. 3). Tama virus, a third virus in the Luteo-Sobemo clade (Fig.2, C), was only detectable by PCR in Japanese samples.

In our *D. suzukii* collections we detected reads from three separate Nora viruses, *D. melanogaster* Nora Virus (Habayeb et al., 2006), *D. immigrans* Nora Virus (van Mierlo et al., 2014) and the new Nora virus, most closely related to that of *D. immigrans,* but sufficiently divergent from both (37.1% and 30.4% amino-acid divergence at the RdRp locus, respectively) to merit description (Fig. 2, D). This clade of viruses also evidently infects other families of ′fruit fly′, as they are detectable in the transcriptomes of two species of tephritids *(Bactrocera latifrons* and *Ceratitis capitata),* and can also be found in the transcriptomes of their parasitoid, *Fopius arisanus* (Fig. 2, D).

Beult virus was the most geographically widespread virus we identified: we detected Beult virus across sampling locations and years, with reads being especially abundant in samples from the UK in 2014 and Japan in 2016. Belonging to a clade of Virga-like viruses (Fig. 2, E), it is very closely related to Bofa virus and Buckhurst virus of *D. melanogaster* and *D. obscura,* respectively (Webster et al., 2016). We identified two different haplotypes of this virus, which share a 98.9% nucleotide similarity: one from the UK, and a second divergent lineage from Japan. Saiwaicho virus (Fig. 2, F), closely related to a group of viruses described as Negeviruses by Vasilakis *et al.* (2013), and Luckshill virus (Fig. 2, G) belonging to a cluster of viruses with close relationship to the Togaviridae, both also fall within the Hepe-Virga clade of +ssRNA viruses. For this clade viruses we were able to identify domains for transferases, helicases, and polymerases (Fig. 3), with the exception of Cyril virus, which was detected from a fragment of the first large virgavirus ORF, encompassing only transferase and helicase domains. Phylogenetic analysis for this virus was therefore performed using the transferase coding sequence (Fig. 2, H).

We detected a single Nido-like virus in our samples from the UK and Japan. We have provisionally named this Fuefuki virus, and it has the longest contig recovered for any of our putative viruses, at over 16.5 kb. Within this near-complete genome we identify five ORFs but only one conserved domain: the RdRp (Fig. 3). Fuefuki virus is very closely related to Wuhan nido-like virus 1 (Shi et al., 2016) at 94.8% amino acid similarity in the polymerase. Along with Hubei *Tetragnatha maxillosa* virus 7 and Wuhan insect virus 19 (Shi et al., 2016) these four viruses form a distinct cluster near to the *Coronaviridae,* a family containing some notable vertebrate pathogens, including the SARS virus (Fig. 2, I).

### Viruses with single-stranded negative sense RNA genomes

Five of the viruses we detected are expected to have ‐ssRNA genomes. Three of these belong to the Bunya-Arena clade of viruses: Notori virus, Ditton virus, and Barming virus (Fig. 4, A–C). Notori and Ditton viruses can be further classified as Phasmaviruses. These were detected in our samples as contigs of around 7kb in length that represent complete, or near-complete L-segments (Bishop and Shope, 1979) (Fig. 5). Barming virus, the third putative Bunya-Arena clade virus we identified, belongs to the Phlebo-like cluster of the clade. It too is known from a contig of just over 6kb, also representing the L-segment of the Bunyavirus genome, consisting of one ORF containing the viral RdRp (Fig. 5). The closest relative of Barming virus was a viral-like sequence identified in the TSA database from *Colletotrichum cereale,* a plant disease that has been found to cause crown rot anthracnose of turf grass (Crouch et al., 2006).

**Figure 4.**
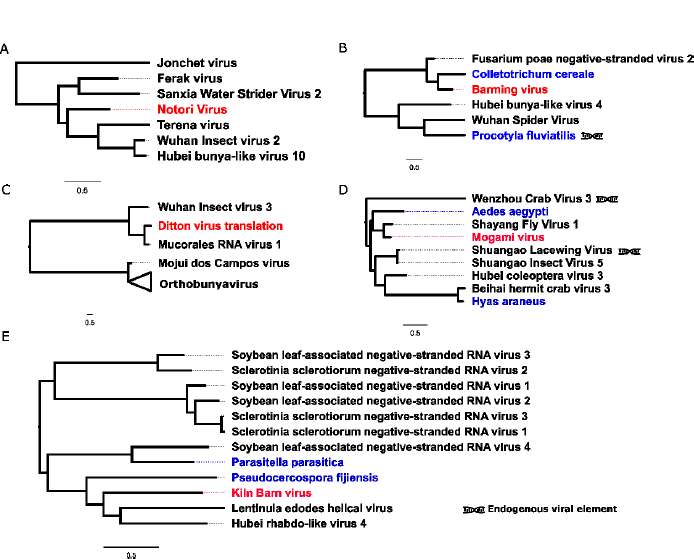
Negative sense single-stranded RNA viruses. Midpoint-rooted, maximum-likelihood trees were inferred from viral polymerase sequences. Scale bar represents 0.5 substitutions per site. Viruses newly described in association with *D. suzukii* (red) are highlighted alongside viral-like sequences identified in public transcriptome datasets (blue). Viruses previously described by the original authors as endogenous viral elements are also marked. Tree A: Viruses close to Phasmaviruses in the Bunya-Arena group; B: Viruses belonging to the Phlebo-like cluster of the Bunya-Arena group; C: Orthobunyaviruses (collapsed) and small sister clade consisting of three viruses, including the newly described Ditton virus; D: Cluster of the Chuviruses; E: Cluster of viruses close to Chuviruses in the Mono-Chu clade. Complete trees are provided in supporting file S3_data.

**Figure 5.**
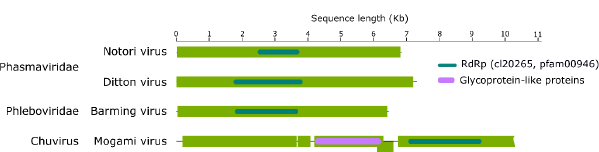
The structure of selected ‐ssRNA virus genomes for which we recover complete or near complete genome sequences. Outer (green) boxes represent boundaries of ORFs and inner boxes represent the relative position of conserved domains identified with reference to NCBI Conserved Domain Database. Waved outer boxes represent incomplete ORFs.

The remaining ‐ssRNA viruses we identified belong to the Mono-Chu clade of ‐ssRNA viruses. From fly samples collected in the UK in 2014 we identified Kiln Barn virus, represented by a 3.7 kb contig containing the RdRp coding domain. The recovery of this segment allowed phylogenetic analysis and the design of primers for RT-PCR detection, however, the remainder of this virus’ genome could not be accurately reassembled and is therefore not annotated in figure 5. Kiln Barn virus clusters phylogenetically with a group of viruses close to the Chuviruses *sensu stricto,* and we find its closest relatives to be Hubei rhabdo-like virus 4 (Shi et al., 2016) and a viral sequence identified in the transcriptome of the Shiitake mushroom fungus *Lentinula edodes* (AGH07920.1).

The other virus we identified from this clade, Mogami virus, is closely related to Shayang fly virus 1, a Chuvirus detected in Chinese Diptera (Shi et al., 2016), and was represented by a 10.5kb contig in which from which we are able to identify both glycoprotein and polymerase ORFs.

### Viruses with double-stranded RNA genomes

We discovered three viruses predicted to possess double-stranded RNA genomes. These included two Totiviruses, Snodland virus and Larkfield virus, both represented by partial protein coding sequences. Both have closest relatives discovered in insect pool sequencing by Shi et al. (2016). Larkfield shares a cluster within the Totiviruses which includes a number of ant viruses: two discovered by Koyama et al. (2015) and Koyama et al. (2016) in genus *Camponotus,* and one found here as a virus-like sequence in a published transcriptome of the black garden ant: *Lasius niger* (Fig. 6). Its closest relative, Hubei toti-like virus 14, is described as an endogenous viral element (Shi et al., 2016). Snodland virus clusters with a small group of other insect viruses, neighbouring a cluster of mycoviruses associated primarily with powdery mildews (Fig. 6).

**Figure 6.**
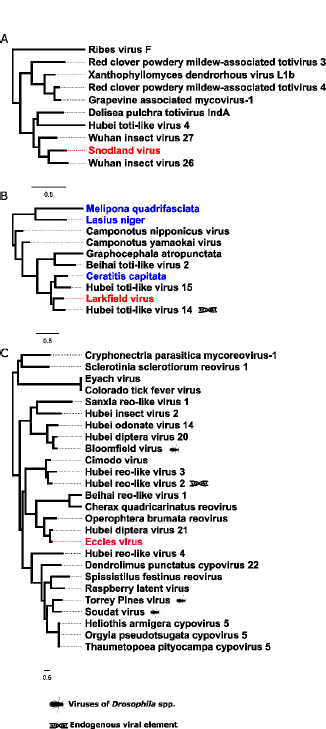
Double stranded RNA viruses. These midpoint-rooted, maximum-likelihood trees were inferred from viral polymerase sequences. Putative viruses newly described in association with *D. suzukii* (red) are highlighted alongside viral-like sequences identified in public transcriptome database (blue). Viruses previously described from a *Drosophila* spp. and viruses described by the original authors as endogenous viral elements are also marked. Tree A: Totiviruses, *Totiviridae;* B: Viruses belonging to a clade of the *Totiviridae,* Toti-Chryso clade; C: Reoviruses, including Coltiviruses (Eyach virus and Colorado tick fever virus) and viruses close to Fijiviruses. Complete trees are provided in supporting file S3_data.

The final dsRNA virus identified, Eccles virus, is our only representative of a virus family that has been previously advocated for the biological control of insect pests (Peng et al., 2000): the *Reoviridae.* Eccles virus is most closely related to Hubei Diptera virus 21 (Shi et al., 2016) and a reovirus of the geometrid, *Operophtera brumata* (Graham et al., 2006). Homology predicts this virus has a multipartite genome consisting of 11 segments, although we were only able to assemble 6 of those segments from our samples.

### Known Drosophila viruses

We also detected 18 further viruses previously described from other species of *Drosophila.* Three known viruses were detected at very high levels (below), and are therefore highly likely to represent infections of *D. suzukii.* The first of these is Brandeis virus (MF953177), the genome of which is reported here for the first time (Fig. 2). Although originally detected by Webster et al. (2015) in public *D. melanogaster* transcriptome datasets (PRJNA159179; Rodriguez et. al., 2012) and provisionally named, it has not previously been detected in wild flies. It is detected here at high levels (26.8% of all remapped virus reads) in *D. suzukii* samples from France in 2013. Brandeis virus belongs to the Hepe-Virga clade of +ssRNA viruses and is closely related to Muthill virus, a virus previously detected in associated with *D. immigrans* (Webster et al., 2016). We were able to assemble a contig of 10.7 kb, which given homology to closely related virga-like viruses is likely to represent a near-complete genome (Fig. 3). The other previously reported *Drosophila* viruses that we reidentified with confidence here are the iflaviruses Kinkell virus and La Jolla virus. Kinkell virus, first described by Webster et al. (2016) was detectable in *D. suzukii* from the UK in 2016, and La Jolla in all samples from all locations. La Jolla virus reads were detected at high abundance in all our samples, comprising up to 30.7% of viral reads in British flies from 2014, and on average 15.0% of virus reads across all samples.

Four viruses of other *Drosophila* species also appear to be present in *D. suzukii* populations. For example, Corseley virus, a virus most associated with *D. subobscura* (Webster et al., 2016), which was detected at fairly high levels in British caught *D. suzukii* from 2016. It is uncommon in other *Drosophila* species (Webster et al., 2016) and is sufficiently divergent from any newly described *D. suzukii* viruses to minimise cross-mapping of reads. Galbut and Chaq viruses are both known infectious agents of *D. melanogaster,* but appear to be at high levels in 2015 *D. suzukii.* Cross-mapping to these viruses is unlikely due to their divergence from other *Drosophila* viruses, and host species contamination is unlikely to explain the high numbers of re-mapped reads observed. An unnamed cripavirus of *D. kikkawai* virus reported by Webster et al. (2015) may represent true association for the same reasons. It was detected at low levels in Japanese flies only. Bloomfield virus, a reovirus of *D. melanogaster,* also likely represents true association with *D. suzukii* as we identified a divergent haplotype of one of the 10 genomic segments in *D. suzukii* that has not previously been seen in *D. melanogaster.* It is tempting to speculate that this reflects a history of host shifting and segment reassortment in this virus.

The remaining previously published viruses, were detected at much lower levels (S3_table). Some of these may represent a low level of cross mapping from newly described but closely related viruses. To test this possibility, we remapped short reads identified as mapping to known *Drosophila* viruses back to their close relatives in *D. suzukii.* This identified two instances where notable cross-mapping between known viruses was possible. The few reads mapping to Prestney Burn virus (Webster et al., 2016) are possibly mismapped Teise virus reads, as 1,189 of the 6,400 reads mapping to Prestney Burns virus also align preferentially to two specific regions of the Teise RNA 1 fragment. Similarly, 27,650 of 90,928 reads mapping to *D. melanogaster* Nora virus also align to the *D. suzukii* Nora virus. In addition, a number of reads may result from sample contamination by misidentified flies and/or library cross-contamination (such as barcode-switching, see: Sinha et al., 2017; Kircher et al., 2011; and Ballenghein et al., 2017). This includes viruses with no close relative associated with *D. suzukii,* such as Thika virus, Craigies Hill virus and Ashworth virus *(unpublished),* or viruses with biologically constrained host ranges, such as the Sigma viruses, along with *Drosophila* A virus (DAV), Drosophila C virus (DCV), and *D. melanogaster* Nora virus that were known to be present in *D. melanogaster* samples run alongside the 2016 *D. suzukii* samples.

### Virus abundance and composition varies among samples

To estimate the amount of virus in each of our samples we mapped all raw reads back to new and previously published putative *Drosophila* virus genomes (Fig. 1). The percentage of non-rRNA reads that mapped to any *Drosophila* virus varied from 0.09% in the poly-A selected French sample up to 5.14% in UK sample from 2016, with an average of 4.27% of reads being viral in Japanese and British pools. Remapping of reads generated by strand specific sequencing (British and Japanese samples), showed that all viruses with negative sense genomes were represented by between 15% and 49% positive sense reads; viruses with double stranded RNA genomes by 53.3% to 70.8% positive reads; and positive sense viruses 88.3% to 100% positive reads (Fig. 7). The only positive sense ssRNA viruses that lacked negative sense reads were represented by less than 2000 reads in total, although in some cases the proportion of negative sense reads was very low. These included Teise virus and La Jolla virus, which displayed extremely large numbers of reads (3,974,042 and 1,326,799 respectively) and the latter of which is a confirmed infectious agent of Drosophila (Webster et al., 2015).

**Figure 7.**
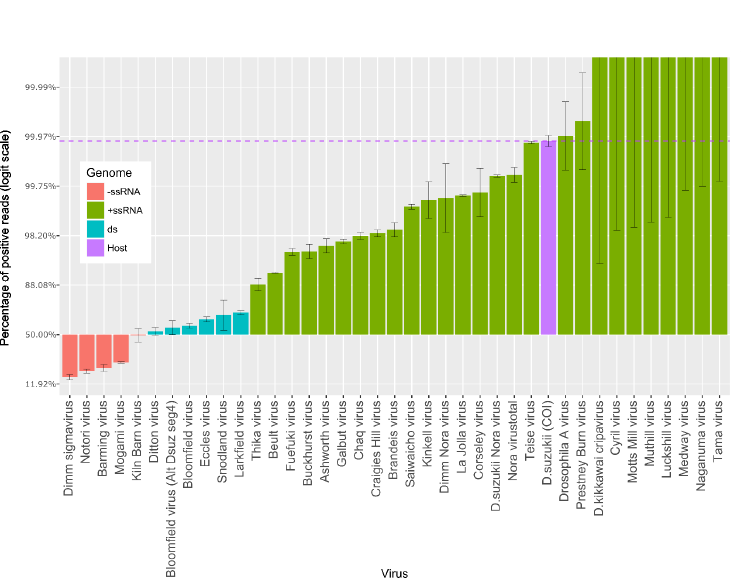
Percentage of reads positive sense remapping to virus or host genomes. Percentages presented on logit scale. Bars represent binomial 95% confidence intervals calculated with logit parameterization. Dashed line represents the percentage of positive reads mapping to *D. suzukii* (Host) COI gene.

The virus composition varied markedly among samples from different times and locations (Fig. 1). Six of the newly described viruses were probably only present in Japanese samples: Mogami virus, Notori virus, Naganuma virus, Saiwaicho virus, Tama virus, and *D. suzukii* Nora virus, whereas many of the new and previously described viruses are found only in the fly′s invasive range and are absent, or at negligible levels, in native, Japanese samples. Despite applying a detection threshold for very low viral read numbers, there are several sources of error when attempting to analyse patterns of virus sharing among years or sampling locations. For example, barcode switching (Sinha et al., 2017, Kircher et al., 2011, Ballenghien et al., 2017) and other sources of cross-contamination between libraries sequenced together on the Illumina platform may allow miss-assignment of reads between the Japanese and British samples from 2016, and also from other drosophilid libraries analysed at the same time. Furthermore, cytochrome oxidase read mapping suggests a small proportion of contaminating reads deriving from *D. melanogaster* and *D. immigrans* were present in some of our datasets. For example, in the Japanese sample of 2016 1.3% of COI reads mapped to *D. immigrans* (potentially misidentified larvae) and in the UK sample of 2015 0.74% of reads mapped to *D. melanogaster.* The *D. melanogaster* reads may represent misidentification or cross-mapping, as the species are quite closely related, but it is more likely that they are the result of contamination across libraries through barcode switching as *D. suzukii* samples were sequenced in parallel with unrelated drosophilid libraries.

## Discussion

Here we make a first survey of the viruses associated with the invasive *Drosophila* pest *D. suzukii* in its native and invasive ranges. Alongside 18 new viruses, not previously described from any organism, we confidently identified a further seven viruses associated with this novel invasive host that had previously been described from other *Drosophila* species. Some novel viruses were detected solely from the native range of *D. suzukii* and others from the invasive range, but rarely from both habitats.

These viruses were identified from metagenomic sequencing of samples of wild *D. suzukii.* Although their presence as RNA but not DNA implies that they are not expressed endogenised viral elements (i.e. EVEs), it remains possible that some are not truly infections in this fly, but may be contaminants of the surface of the fly or infect a commensal, pathogen, or food organism within the fly′s gut lumen. However, we believe that this is unlikely to be the case for most sequences, as previous studies that additionally used the presence of virus-derived 21nt short interfering RNAs to demonstrate active replication (Webster et al., 2015) found that the majority of viruses identified in similar metatransciptomic sequencing of *D. melanogaster* constituted active infections. For most of these viruses active replication is further supported by the relative proportions of positive and negative sense reads mapping to each virus. Although the exact ratio of positive to negative strand RNA is known to fluctuate through the course of infection (Martínez et al., 2011; Thébaud et al., 2009), all viral read counts deviated from the ratio expected if no replication was occurring (Fig 7). This was unambiguous for all of the ‐ssRNA viruses and dsRNA viruses, which showed substantial numbers of the positive sense sequences required for protein synthesis and replication, and strongly supportive for most +ssRNA viruses, almost all of which displayed some of the negative sense reads expected from replication intermediates. There is also a possibility that cross-species contamination, barcode-switching and cross-mapping could result in spurious host allocation, but this is not compatible with the read number or distribution of reads for the majority of viruses (above).

In addition, recent large-scale invertebrate virus discovery projects (Shi et al., 2016) give us a greatly increased confidence in the phylogenetic relationships of newly identified virus sequences. In particular, although some virus taxa have a diverse host range, it seems reasonable to infer that *D. suzukii* is the true host for viruses with very close relatives confirmed to infect another insect. For example, Mogami virus (Chuvirus) is distantly related to any known *Drosophila* virus, but is closely related to Shayang Fly virus 1 (Shi et al., 2016) and clusters within a group of viruses that are only described from insect samples (see Fig. 4, D). Nevertheless, this pattern is not true for all viruses described here. Specifically, two of the 18 novel viruses in this study (Ditton virus and Barming virus), are more closely related to Mycoviruses than they are to any entomopathogenic viruses and one (Luckshill virus) is most closely related to a sequence found in a parasitic nematode of ruminants. And, while this pattern does not exclude the possibility of these being true viruses of *D. suzukii—*as many viral families contain a broad range of hosts including those of different phyla and patterns of host switching are still little understood—these are among the best candidates to be infections of *Drosophila* parasites or gut fauna, rather than *D. suzukii* itself.

The potential for these viruses to be used as biological control agents is currently unclear. Commercially successful viral biocontrol agents have in the past only come from the dsDNA virus family *Baculoviridae,* which was not represented in our collections, and most lineages represented here have not been investigated for their ability to be cultured and applied as control agents. Indeed, few viruses in the identified families have been successfully isolated for experimentation, and many are known only from metagenomic sequencing. The only virus family we found in associated with *D. suzukii* that has any history as a control agent (Zeddam et al., 2003, Peng et al., 1998, Peng et al., 2000) is the reovirus ′Eccles virus′. Eccles virus was relatively rare in our samples, but this may speak to the potential pathogenicity of the virus, as flies harbouring a particularly pathogenic virus, especially one that has a short latency period, may be less likely to visit baited traps (Gupta et al., 2017). Further investigation of this virus, including isolation and pathogenicity assays, are needed before any further conclusions can be drawn about its utility as a control agent. Viruses potentially lethal to *D. suzukii* may also await discovery in other species of *Drosophila.* Indeed, pathogens have the potential to display increased virulence following a host shift event (Longdon et al., 2015) and the susceptibility of *D. suzukii* to viruses of *D. melanogaster* has been shown experimentally (Cattel et al., 2016, Lee and Vilcinskas, 2017). Here we show the potential association of viruses from *D. melanogaster, D. immigrans* and *D. subobscura* with *D. suzukii* in the wild. Further investigation of the viral community experienced by many different *Drosophila* in nature may, therefore, be of both academic and applied interest.

Given our focus on an invasive species, the potential for a shift in the virological environment associated with invasion is of particular interest. Theory predicts that organisms may experience a ′release′ from natural enemies, including pathogens, in their invasive range due to low host densities and founder effects at the invasive edge (Keane and Crawley, 2002): However, this idea remains contentious, as supporting evidence is limited (Colautti et al., 2004). It has also been hypothesised that invasives, rather than experience a drop in overall number of enemies, undergo a shift in the type of enemy encountered, from co-evolved specialists in the native range to more generalist enemies, quickly able to adapt to a new host, in the naturalized range (Joshi and Vrieling, 2005). In this study, we do detect an apparently marked difference in the virus communities of flies from different areas within its expanding geographical range. Although the potential for cross-mapping, and a low level of species contamination, mean that these findings should be treated with some caution, five of the new viruses described (Saiwaicho virus, Tama virus, Mogami virus, Naganuma virus and Notori virus) were only detected at high levels in Japanese (native) flies. These five viruses are not particularly closely related to any previously described *Drosophila* viruses (Fig. 2 and Fig. 4) and may represent a more specialized relationship with *D. suzukii.* In contrast, the three most ubiquitous viruses across all samples, La Jolla virus, Teise virus and Beult virus are either a known generalist (La Jolla) or very closely related to a virus in another related hosts (Fig.2, A and E). If confirmed, this pattern could reflect a shift in natural enemy type from native to invasive range of *D. suzukii.*

## Acknowledgements

We thank Michelle Fountain, Bethan Shaw, Adrian Harris, Maddie Cannon, and the entomological team at East Malling research for support in collecting British flies. We thank members of the Nakai lab at the Tokyo University of Agriculture and Technology for making us welcome, and the staff at the TUAT Field Museum Tama hills, Yamagata Prefectural Agricultural Research Centre, Agriculture Centre Kaju Institute, Fukushima and Yamanashi Prefecture Outside Organization Fruit Tree Test Site for support during field collections. We thank Edinburgh Genomics for sequencing, and members of the Obbard and Vale labs for discussion. This work was funded by a Wellcome Trust Research Career Development Fellowship (WT085064) to DJO and an Agricultural and Horticultural Development Board grant to DJO and JVC to support NCM′s PhD. Work in DJO′s laboratory was partly supported by a Wellcome Trust strategic award to the Centre for Immunity, Infection, and Evolution (WT095831).

## Data availability

All raw unannotated contigs are provided in supporting file S1_data (DOI: 10.6084/m9.figshare.5649829); all alignments are provided in supplementary material S2_data (DOI: 10.6084/m9.figshare.5650117); all large phylogenetic trees from which figures are taken are available in S3_data (DOI: 10.6084/m9.figshare.5650132); a table of known *Drosophila* viruses detected in *D. suzukii* is available in S1_table (DOI: 10.6084/m9.figshare.5650147); a table of PCR primers used for virus detection is in S2_table (DOI: 10.6084/m9.figshare.5650156); Number of reads mapping to all known drosophila viruses and to novel D. suzukii viruses is in S3_table (DOI: 10.6084/m9.figshare.5830644). Coverage depth graphs for novel virus genomes are shown in S1_figure (DOI: 10.6084/m9.figshare.5893324).

All raw reads have been submitted to the NCBI sequence read archive under project accession PRJNA402011 (Japan SRR6019484; France SRR6019487; Kent: SRR6019485, SRR6019486, and SRR6019488). All novel virus genomes are submitted separately to the NCBI sequence read archive under the accessions outlined in table 1.

